## Supplemental figures and tables for "A single, improbable B cell receptor mutation confers potent neutralization against cytomegalovirus"

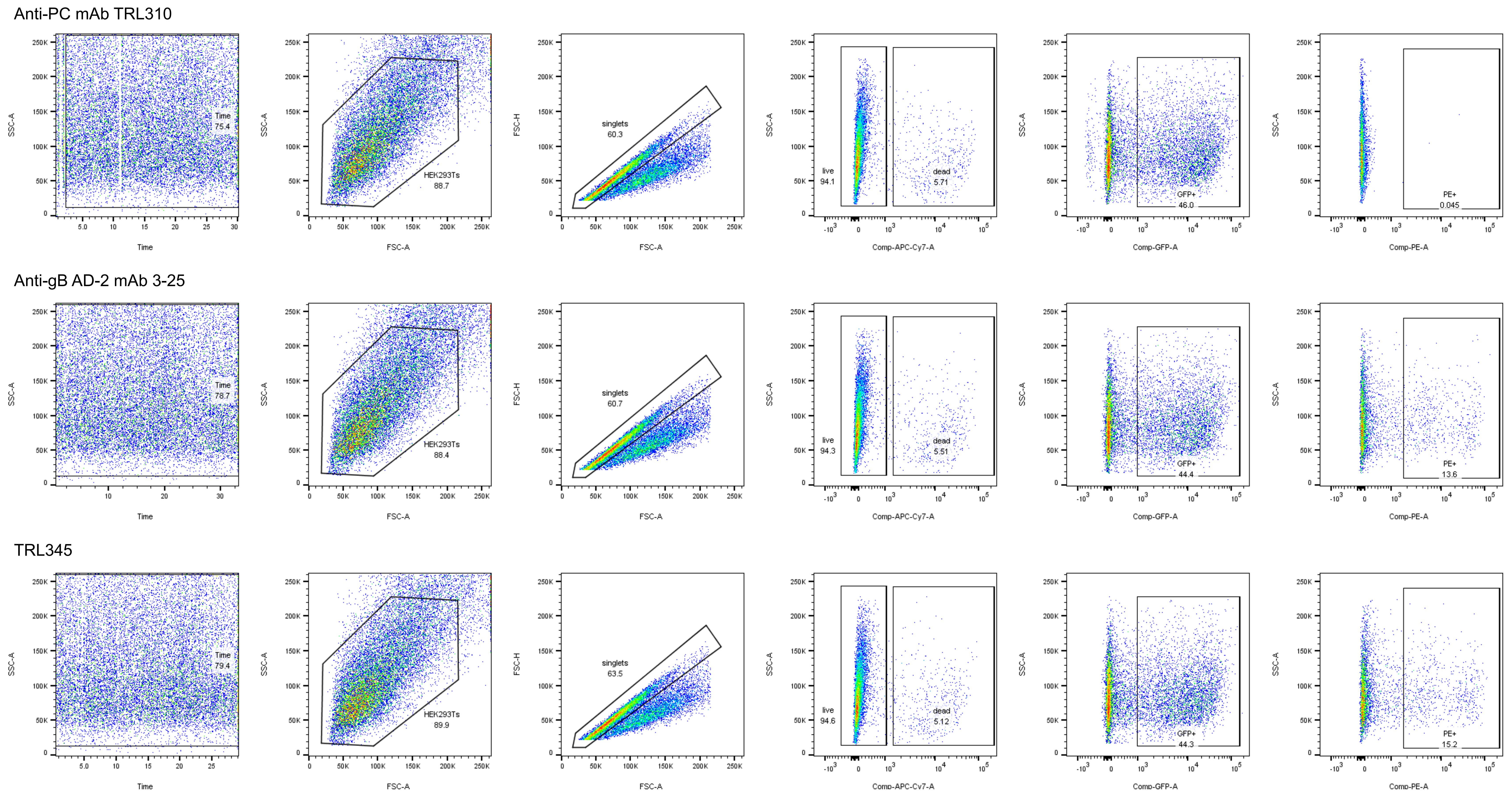

**Fig. S1. Gating strategy to determine the % of binding to cell-associated gB.** We co-transfected DNA plasmids that separately expressed full-length gB (Towne strain) or GFP. We coincubated gB and GFP-expressing cells with mAbs in a three-point, 10-fold serial mAb dilution, then detected anti-gB AD-2S1 mAb binding with PE-conjugated anti-human IgG Fc. We defined the minimum threshold of positive expression as binding by an anti-CMV pentameric complex mAb “TRL310” and reported the % of GFP-expressing, live HEK293T singlets as “% gB-transfected cell binding.”

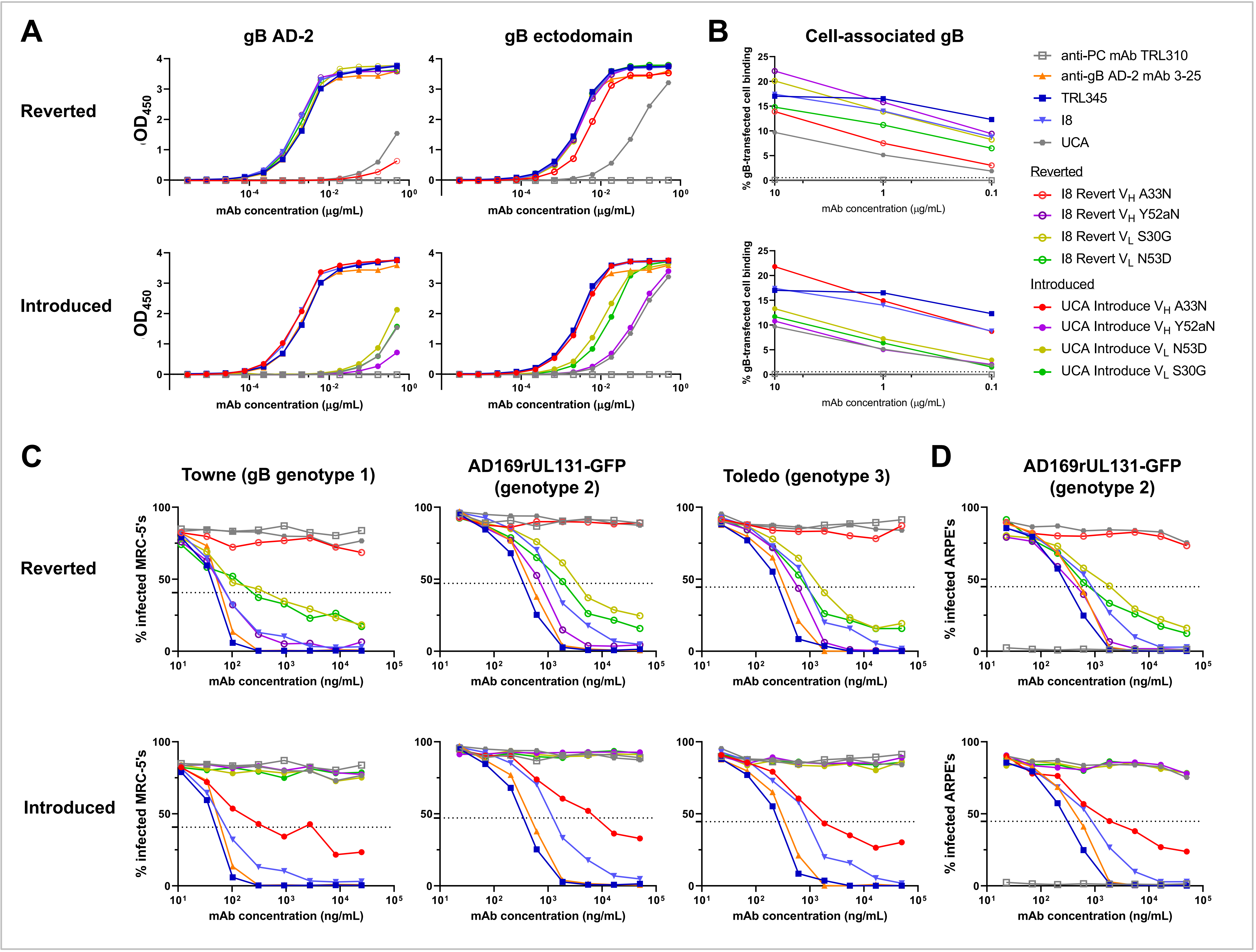

**Fig. S2. The VH A33N mutation was both necessary and sufficient for high gB AD-2 binding, binding to cell-associated gB, and neutralization function.** We produced the 17 clonally related mAbs of the TRL345 lineage and measured the following:

(A) Binding magnitude to gB AD-21 peptide and gB ectodomain by ELISA.
(B) Binding to cell-associated gB. Binding of mAbs was determined by coincubating mAbs in a serial dilution with HEK293T epithelial cells-transfected with full-length gB and GFP. The %binding was calculated as the % of GFP-expressing cells bound by the anti-gB AD-2 mAb, detected by flow cytometry.
(C) Neutralization function against CMV strains Towne, AD169rUL131-GFP (AD169r), and Toledo on MRC-5 fibroblasts.
(D) Neutralization function against CMV strain AD169rUL131-GFP on ARPE epithelial cells.

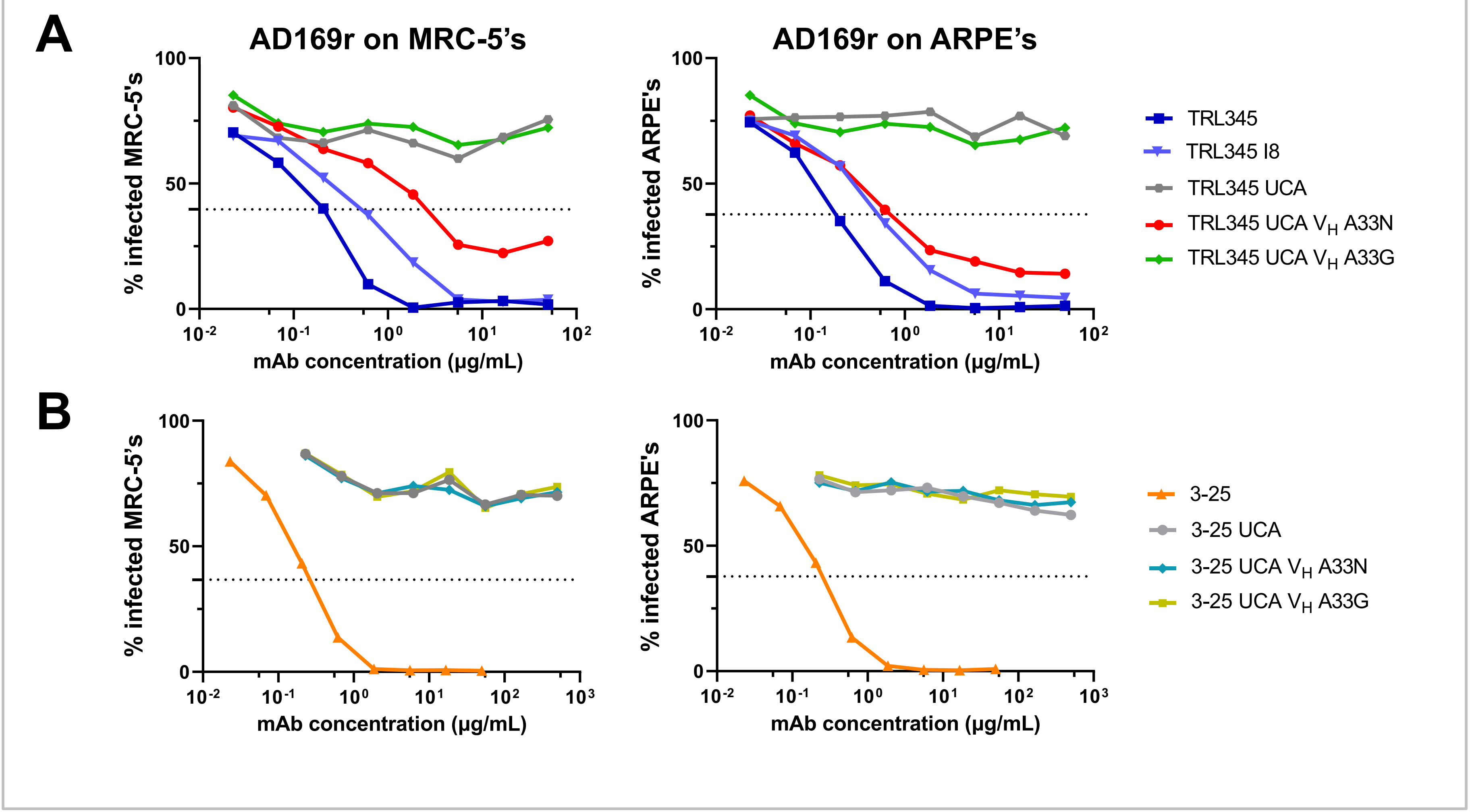

**Fig. S3. Introduction of the $V_H$ A33G mutation to the UCA of either the TRL345 or 3-25 lineages did not confer neutralizing function.** We measured neutralization of the CMV strain AD169rUL131-GFP on MRC-5 fibroblasts or ARPE epithelial cells for the following mAbs:

(A) TRL345 lineage antibodies and TRL345 UCA with either the $V_H$ A33N or A33G mutations.

(B) 3-25 lineage antibodies and 3-25 UCA with either the $V_H$ A33N or A33G mutations. 3-25 mutant mAbs were coincubated with virus at a 10-fold higher starting concentration than all other mAbs, but even at these higher concentrations, these mAbs were non-neutralizing.

**A** Heavy chains

```
                                                   33                          52a
                                              CDR1                        CDR2
TRL345 UCA    QVQLVESGGGVVQPGRSLRLSCAASGFTFSSYAMHWVRQAPGKGLEWVAVISYDGSNKYYA     60
I8            QVQLVESGGGVVQPGRSLRLSCAASGFTFSSYNMHWVRQAPGKGLEWVAVISNDGSNKYYA     60
TRL345        QVQLVESGGGVVQPGRSLRLSCAASGFTFSDYNMHWVRQAPGKGLEWVAVISIDGTYKYSA     60
3-25 UCA      QVQLVESGGGVVQPGRSLRLSCAASGFTFSSYAMHWVRQAPGKGLEWVAVISYDGSNKYYA     60
3-25          QVQLVESGGGVVQPGRSLRLSCAASGFTFSNHGLHWVRQPPGKGLEWVAVVSKDGTNEHYA     60
                                                 CDR3
TRL345 UCA    DSVKGRFTISRDNSKNTLYLQMNSLRAEDTAVYYCARDGRSVGG--FSGILDPWGQGTLVT    110
I8            DSVKGRFTISRDNSKNTLYLQMNSLRAEDTAVYYCARDGRSVGG--FSGILDPWGQGTLVT    110
TRL345        DSVAGRFSLSRDNSKNTLYLQMNSLRPDDTAIYYCARDGRSVGG--FSGILDPWGQGTLVT    110
3-25 UCA      DSVKGRFTISRDNSKNTLYLQMNSLRAEDTAVYYCAREGYCSGGSCYSGQPDYWGQGTLVT    110
3-25          DSVRGRFTISRDNSKNTLYLLMKSLRLEDTAVYYCAREGYCGDDRCYSGQPDYWGQGILVT    110

TRL345 UCA    VSS     113
I8            VSS     113
TRL345        VSS     113
3-25 UCA      VSS     113
3-25          VSS     113
```

TRL345 I8 binding sites to gB AD-2S1

3-25 binding sites to gB AD-2S1

Early, improbable mutations

**B** Light chains

```
                                            30                          53
                                         CDR1                      CDR2
TRL345 UCA    EIVLTQSPATLSLSPGERATLSCRASQSVSSYLAWYQQKPGQAPRLLIYDASNRATGIPA     60
I8            EIVLTQSPATLSLSPGERATLSCRASQSVGSYLAWYQQKPGQAPRLLIYDASDRATGIPA     60
TRL345        EIVMTQSPATLSLSPGDRATLSCRASQSVGSYLAWYQQKPGQAPRLLMYDSSVRATGIPA     60
3-25 UCA      EIVLTQSPATLSLSPGERATLSCRASQSVSSYLAWYQQKPGQAPRLLIYDASNRATGIPA     60
3-25          EIVLTQFPATLSLSPGERATLSCRASQSVGRYLAWYQQKPGQAPRLLIYDSSNRATGVPA     60
                                               CDR3
TRL345 UCA    RFSGSGSGTDFTLTISSLEPEDFAVYYCQQRSNWPPLTFGGGTKVEIK     107
I8            RFSGSGSGTDFTLTISSLEPEDFAVYYCQQRSNWPPLTFGGGTKVEIK     107
TRL345        RFSGSGSGTDFTLTISSLEPEDFAVYYCQQRNNWPPLTFGGGTKVEIK     107
3-25 UCA      RFSGSGSGTDFTLTISSLEPEDFAVYYCQQRSNWPPLTFGGGTKVEIK     107
3-25          RFSGSGSGTDFTLSISSLEPEDFAVYFCQQRSHWPPLTFGGGTKVEIK     107
```

**Fig. S4. Sequence alignment of anti-gB AD-2S1 mAbs from the TRL345 and 3-25 lineages.** The TRL345 UCA, I8, and mature mAb were aligned with the 3-25 UCA and mature mAb for the (**A**) heavy chains and (**B**) light chains. Amino acid numbering is in the Kabat scheme. CDR regions are underlined. Blue boxes indicate the contact sites for the TRL345 I8 antibody to gB AD-2S1 peptide by crystal structure, and the yellow boxes indicate the contact sites for the 3-25 mature antibody to gB AD-2S1. Red letters indicate amino acid mutations from the respective UCAs of each lineage.

**Table S1. Compiled binding and neutralization responses for TRL345 lineage mAbs and 3-25 mature mAb.**

| ***Assay*** | ***ELISA*** | | ***SPR*** | | | | | | | | ***Cell-associated gB binding*** | ***Neutralization on fibroblasts*** | | | | ***Neutralization on epithelial cells*** |
|---|---|---|---|---|---|---|---|---|---|---|---|---|---|---|---|---|
| | *gB AD-2* | *gB ectodomain* | *gB AD-2* | | | | *gB ectodomain* | | | | *Area under the curve (AUC) for % gB-transfected cell binding* | | | | | |
| ***mAb*** | *$EC_{50}$ (ng/mL)* | *$EC_{50}$ (ng/mL)* | *$k_a$ ($10^6$/M*s)* | *$k_d$ (1/ks)* | *$K_d$ (nM)* | *Chi² (RU²)* | *$k_a$ ($10^6$/M*s)* | *$k_d$ (1/ks)* | *$K_d$ (nM)* | *Chi² (RU²)* | | *Towne $IC_{50}$ (µg/mL)* | *AD169rUL131-GFP $IC_{50}$ (µg/mL)* | *Toledo $IC_{50}$ (µg/mL)* | *Average $IC_{50}$ (µg/mL)* | *AD169rUL131-GFP $IC_{50}$ (µg/mL)* |
| ***TRL345*** | *3.3* | *4.4* | *3.74* | *5.88* | *0.29* | *0.61* | *12.90* | *0.025* | *0.81* | *0.59* | *151.6* | *0.21* | *0.24* | *0.19* | *0.21* | *0.3* |
| ***MAB343*** | *8.0* | *9.6* | *2.24* | *0.64* | *1.57* | *1.02* | *16.00* | *0.013* | *1.92* | *1.29* | *161.1* | *0.35* | *0.23* | *0.22* | *0.27* | *0.23* |
| ***I1*** | *2.6* | *4.0* | *2.61* | *4.16* | *1.59* | *1.46* | *14.10* | *0.013* | *0.92* | *0.72* | *142.7* | *0.4* | *0.31* | *0.25* | *0.32* | *0.4* |
| ***MAB309 (I2)*** | *3.3* | *5.5* | *2.08* | *7.84* | *3.77* | *2.10* | *12.60* | *<0.01* | *0.79* | *0.83* | *124.3* | *0.79* | *0.73* | *0.64* | *0.72* | *1.02* |
| ***MAB318*** | *2.2* | *3.6* | *1.75* | *6.44* | *3.68* | *1.24* | *11.90* | *<0.01* | *0.84* | *0.83* | *132.6* | *0.63* | *0.58* | *0.49* | *0.57* | *0.63* |
| ***MAB310 (I3)*** | *2.3* | *3.8* | *6.36* | *19.80* | *3.11* | *2.43* | *13.50* | *<0.01* | *0.74* | *0.67* | *187.1* | *0.31* | *0.23* | *0.22* | *0.25* | *0.31* |
| ***I4*** | *2.5* | *4.6* | *3.07* | *5.68* | *1.85* | *1.49* | *15.30* | *0.011* | *0.71* | *1.20* | *129.7* | *0.37* | *0.3* | *0.24* | *0.31* | *0.36* |
| ***MAB338*** | *3.4* | *3.9* | *2.15* | *4.27* | *1.99* | *1.10* | *13.60* | *<0.01* | *0.74* | *1.01* | *136.3* | *0.36* | *0.32* | *0.32* | *0.33* | *0.36* |
| ***MAB319*** | *6.5* | *4.0* | *2.25* | *2.03* | *0.90* | *0.69* | *11.80* | *<0.01* | *0.85* | *0.95* | *130* | *0.62* | *0.64* | *0.57* | *0.61* | *0.78* |
| ***I5*** | *5.2* | *4.4* | *3.37* | *18.60* | *5.52* | *0.08* | *13.90* | *<0.01* | *0.72* | *1.20* | *122.6* | *0.52* | *0.46* | *0.41* | *0.46* | *0.59* |
| ***MAB313*** | *2.2* | *3.5* | *11.7* | *51.20* | *4.38* | *3.35* | *13.10* | *0.011* | *0.82* | *0.88* | *154.5* | *0.26* | *0.18* | *0.18* | *0.21* | *0.2* |
| ***MAB316*** | *2.5* | *3.7* | *2.78* | *8.23* | *2.96* | *1.30* | *12.60* | *<0.01* | *0.79* | *0.93* | *163* | *0.46* | *0.42* | *0.37* | *0.42* | *0.56* |
| ***I6*** | *2.9* | *4.2* | *2.94* | *7.92* | *2.69* | *1.68* | *14.50* | *<0.01* | *0.69* | *1.21* | *160.7* | *0.35* | *0.31* | *0.28* | *0.31* | *0.36* |
| ***I7*** | *2.5* | *4.1* | *3.09* | *13.80* | *4.47* | *0.84* | *10.90* | *<0.01* | *0.92* | *1.03* | *146.4* | *0.49* | *0.39* | *0.3* | *0.39* | *0.46* |
| ***I8*** | *2.5* | *4.6* | *8.56* | *199.00* | *23.20* | *4.59* | *9.95* | *<0.01* | *1.01* | *1.23* | *132.6* | *2.78* | *0.83* | *0.83* | *1.48* | *0.79* |
| ***UCA*** | *377.1* | *171.7* | *6.15* | *170.00* | *27.60* | *0.04* | *8.27* | *0.264* | *31.90* | *0.74* | *97.3* | ***50*** | ***50*** | ***50*** | ***50*** | ***50*** |
| ***3-25*** | *3.0* | *3.7* | *6.11* | *27.00* | *4.42* | *2.28* | *12.30* | *<0.01* | *0.81* | *0.53* | *148.9* | *0.27* | *0.31* | *0.26* | *0.28* | *0.48* |

*In the neutralization assays, mAbs were run in an 8-point 3x dilution series, starting at 50 µg/mL. mAbs that did not achieve 50% inhibition of maximal infection were determined to be "non-neutralizing," and their $IC_{50}$ values were set to 50.0 µg/mL (light gray boxes), which was the maximal concentration measured. The positive control mAb 3-25 was assessed in all assays (dark grey boxes).*

**Table S2. Mutational probability analysis using ARMADiLLO algorithm.**

| | *Heavy chain* | | | *Light chain* | | | *mAb total* |
|---|---|---|---|---|---|---|---|
| ***mAb*** | *# nt mutations* | *# AA mutations* | *# AA mutations (Pr<0.02)* | *# nt mutations* | *# AA mutations* | *# AA mutations (Pr<0.02)* | *# AA Mutations (P<0.02)* |
| ***MAB345*** | *30* | *12* | *6* | *7* | *12* | *2* | *8* |
| ***MAB343*** | *21* | *13* | *5* | *4* | *8* | *3* | *8* |
| ***I1*** | *16* | *10* | *5* | *4* | *7* | *3* | *8* |
| ***MAB309 (I2)*** | *25* | *18* | *7* | *8* | *16* | *2* | *9* |
| ***MAB318*** | *25* | *18* | *8* | *9* | *18* | *2* | *10* |
| ***MAB310 (I3)*** | *23* | *15* | *6* | *7* | *12* | *2* | *8* |
| ***I4*** | *16* | *10* | *5* | *3* | *6* | *2* | *7* |
| ***MAB338*** | *40* | *19* | *6* | *6* | *16* | *4* | *10* |
| ***MAB319*** | *30* | *14* | *5* | *4* | *9* | *2* | *7* |
| ***I5*** | *21* | *10* | *5* | *3* | *6* | *2* | *7* |
| ***MAB313*** | *38* | *21* | *8* | *8* | *16* | *3* | *11* |
| ***MAB316*** | *35* | *20* | *6* | *7* | *14* | *5* | *11* |
| ***I6*** | *30* | *17* | *6* | *6* | *13* | *4* | *10* |
| ***I7*** | *12* | *5* | *3* | *2* | *4* | *2* | *5* |
| ***I8*** | *5* | *2* | *2* | *2* | *3* | *2* | *4* |
| ***UCA*** | *0* | *0* | *0* | *0* | *0* | *0* | *0* |

**Table S3. X-ray crystallographic data collection and refinement statistics.**

| | **I8 Fab + gB AD-2S1** |
|---|---|
| **PDB ID** | **----** |
| **Data collection** | |
| Space group | *P 21 21 21* |
| Cell dimensions | |
| *a, b, c* (Å) | 66.9, 74.7, 104.9 |
| α, β, $\gamma$ (°) | 90.0, 90.0, 90.0 |
| Wavelength (Å) | 0.9792 |
| Resolution (Å) | 66.88-1.80 (1.84-1.80) |
| Unique reflections | 48,200 (2,826) |
| $R_{merge}$ | 0.061 (0.130) |
| $R_{pim}$ | 0.040 (0.087) |
| I/σI | 17.2 (9.6) |
| $CC_{1/2}$ | 0.997 (0.982) |
| Completeness (%) | 97.5 (97.7) |
| Multiplicity | 6.0 (6.0) |
| Wilson *B*-factors (Å$^2$) | 13.4 |
| **Refinement** | |
| Resolution | 56.39-1.80 (1.84-1.80) |
| Unique reflections | 48,141 (2,634) |
| $R_{work}/R_{free}$ (%) | 15.0/17.4 (17.9/20.7) |
| No. atoms | |
| Protein | 3361 |
| Water | 723 |
| Ligands | 6 |
| B-factors (Å$^2$) | |
| Protein | 15.3 |
| Water | 31.7 |
| Ligands | 14.0 |
| R.m.s. deviations | |
| Bond lengths (Å) | 0.01 |
| Bond angles (°) | 0.77 |
| Ramachandran | |
| Favored (%) | 98.6 |
| Allowed (%) | 1.4 |
| Outliers (%) | 0.0 |
